# Evolutionary successful strategies in a transparent iterated Prisoner’s Dilemma

**DOI:** 10.1101/524348

**Authors:** Anton M. Unakafov, Thomas Schultze, Igor Kagan, Sebastian Moeller, Alexander Gail, Stefan Treue, Stephan Eule, Fred Wolf

**Affiliations:** Georg-Elias-Mueller-Institute of Psychology, University of Goettingen, Gosslerstrasse 14, 37073, Göttingen, Germany; Max Planck Institute for Dynamics and Self-Organization, Am Fassberg 17, 37077, Göttingen, Germany; Leibniz ScienceCampus Primate Cognition, Kellnerweg 4, 37077, Göttingen, Germany; German Primate Center – Leibniz Institute for Primate Research, Kellnerweg 4, 37077, Göttingen, Germany; Campus Institute for Dynamics of Biological Networks, Hermann Rein Strasse 3, 37075 Göttingen, Germany; Max Planck Institute for Experimental Medicine, Hermann Rein Strasse 3, 37075 Göttingen, Germany; Bernstein Center for Computational Neuroscience, Am Fassberg 17, 37077, Göttingen, Germany

**Keywords:** Evolutionary game theory, Iterated Prisoner’s dilemma, Transparent games

## Abstract

A Transparent game is a game-theoretic setting that takes action visibility into account. In each round, depending on the relative timing of their actions, players have a certain probability to see their partner’s choice before making their own decision. This probability is determined by the level of transparency. At the two extremes, a game with zero transparency is equivalent to the classical simultaneous game, and a game with maximal transparency corresponds to a sequential game. Despite the prevalence of intermediate transparency in many everyday interactions such scenarios have not been sufficiently studied. Here we consider a transparent iterated Prisoner’s dilemma (iPD) and use evolutionary simulations to investigate how and why the success of various strategies changes with the level of transparency. We demonstrate that non-zero transparency greatly reduces the set of successful memory-one strategies compared to the simultaneous iPD. For low and moderate transparency the classical “Win – Stay, Lose – Shift” (WSLS) strategy is the only evolutionary successful strategy. For high transparency all strategies are evolutionary unstable in the sense that they can be easily counteracted, and, finally, for maximal transparency a novel “Leader-Follower” strategy outperforms WSLS. Our results provide a partial explanation for the fact that the strategies proposed for the simultaneous iPD are rarely observed in nature, where high levels of transparency are common.

## 1 Introduction

Game theory is widely used to account for strategic decision-making in rational agents. Classical game theory assumes that players act either sequentially or simultaneously. Yet, for social behavior these assumptions are often not fulfilled since humans and animals rarely act strictly simultaneously or sequentially. Instead, social agents try to observe their partners and adversaries, and use the others’ behavior to adjust their own actions accordingly [1,2]. Recently a new game-theoretic setting of “transparent games” has been introduced, taking into account action visibility and providing a more realistic model of interactions under time constraints [3]. In transparent games, each player has a certain probability to observe the partner’s choice before deciding on its own action. This probability is determined by the action times of the players. For instance, if Player 1 always acts well before Player 2, the probability to see the partner’s choice is zero for Player 1 and one for Player 2 (corresponds to strictly sequential playing). If both players act approximately at the same time, on average, they have equal probabilities *p*_see_ to see each other’s choices. This probability can range from *p*_see_ = 0 (players cannot take choices of each other into account; this case corresponds to the classical simultaneous game) to *p*_see_ = 0.5 (in every round typically one of the two players sees the choice of the partner and can adjust its own decision). The probability that neither player sees the choice of the partner is equal to 1 — 2 *p*_see_.

One may expect that action visibility would increase cooperation in non-zero-sum games, such as the iterated Prisoner’s dilemma (iPD). However, evolutionary simulations in [3] show that this is not necessarily the case. Evolutionary agents successfully establish cooperation in the iPD for low and moderate transparency by using the classic “Win-stay, lose – shift” (WSLS) strategy [4]. However, for high transparency cooperation drastically decreases (see Fig. 1), and the most frequent strategy is “Leader-Follower”, which does not rely on mutual cooperation [3].

**Fig. 1.**
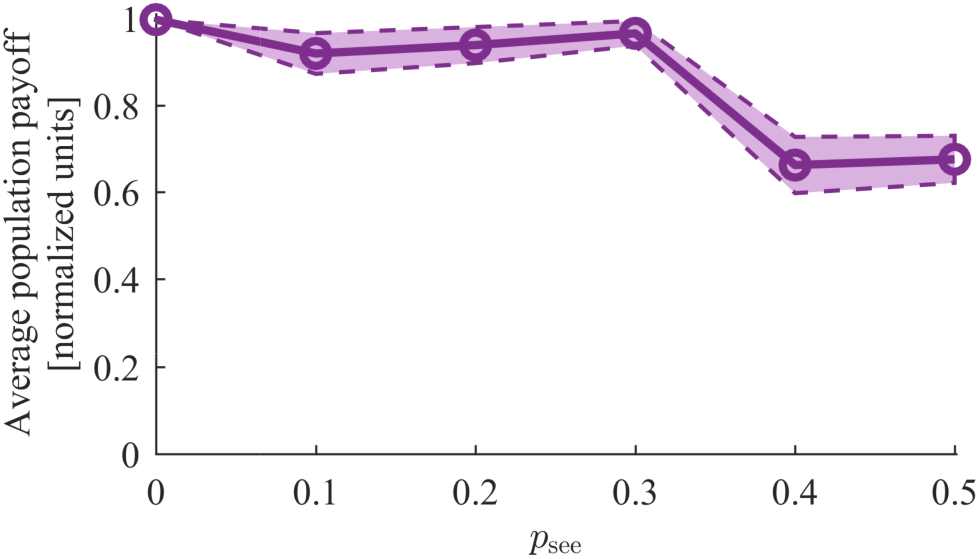
Average fraction of maximal possible payoff (with 95% confidence interval) achieved by the evolutionary evolved population of players in the transparent iterated Prisoner’s Dilemma. Payoff values are computed over 80 runs for probabilities *P*_see_ = 0.0, 0.1,…, 0.5 of a player to see the choice of a partner. Maximal payoff corresponds to mutual cooperation. Stable cooperation was established for low and moderate transparencies, but it was disrupted for *p*_see_ > 0.3 resulting in a significant decrease of average payoff.

This unexpected drop of cooperation in an iPD with high transparency raises new questions. What explains the success of WSLS for the transparent iPD? Why does the Leader-Follower strategy become more frequent for high transparency? Is its success evolutionary stable or is this strategy just transiently successful? To answer these questions, we use evolutionary simulations, since analytic considerations for the transparent games require solving differential equations with many variables. Since most results for simultaneous and sequential versions of the iPD were obtained for strategies taking into account outcomes of only the last interaction (“memory-one strategies”), we also focus on such strategies. We show that the rather complex strategy dynamics associated with *p*_see_ = 0 (simultaneous iPD) [5] is greatly simplified for the transparent settings with *p*_see_ > 0. WSLS is the only non-transient strategy for 0 < *p*_see_ < 0.5. For *p*_see_ > 0.35 all memory-one strategies are evolutionary unstable and replace each other in rapid succession, though WSLS and Leader-Follower are more stable than the others.

Our results complement the findings in [3] by providing a detailed analysis of the evolutionarily successful strategies for the transparent iPD. In particular, we emphasize the superiority of WSLS over other memory-one strategies for most transparency levels. Yet, the fact that WSLS and other strategies developed for the simultaneous iPD are unstable for an iPD with high transparency may partially explain why these classic strategies are rarely observed in nature [6, 7], where high levels of transparency are prevalent [1].

## 2 Methods

### 2.1 Transparent iterated Prisoner’s dilemma

Prisoner’s dilemma [8] is perhaps the most studied game between two players. We refer to [9] for a review. The payoff matrix for this game is presented in Fig. 2. Apparently, the best choice for both players is mutual cooperation, but defection dominates cooperation (i.e., it yields a higher payoff for each individual player regardless of the partner’s choice). Thus any self-interested player would defect (D), although mutual defection results in lowest possible joint payoff. This “paradox” of Prisoner’s dilemma raises a question: under what conditions is it rational to cooperate (C)?

**Fig. 2.**
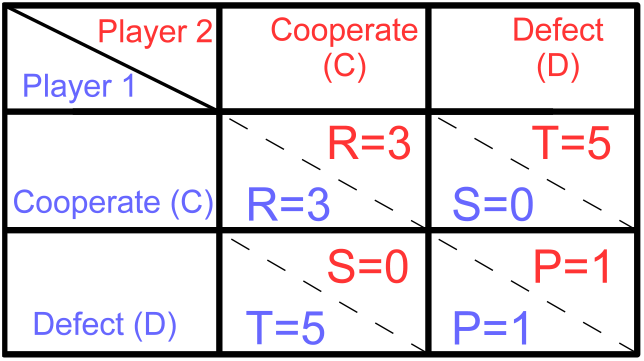
Payoff matrix for Prisoner’s Dilemma. In this game two players adopt roles of prisoners suspected of committing a crime. Each can either betray the other (Defect), or Cooperate with the partner by remaining silent. The maximal charge is five years in prison, and the payoff matrix represents the number of years deducted from it. For example, if Player 1 defects and Player 2 cooperates, then Player 1 gets payoff *T* = 5 and goes free, while Player 2 gets *S* = 0 and has to spend five years in prison. If both prisoners defect, each gets a slightly reduced sentence of four years (*P* = 1) Finally, with mutual cooperation between the prisoners there is only circumstantial evidence left, which is sufficient to sentence both prisoners to two years in prison (*R* = 3).

One possible answer is provided by playing the game repeatedly: in the iPD cooperation between self-interested players becomes plausible, since they can take into account past outcomes but also have to consider that they will play against each other again in future. When experience shows the partner to be trustworthy, cooperation in repeated game becomes a viable option. Traditionally each round of the iPD is considered to be a simultaneous game [8], where players make choices independently. In the recently suggested transparent iPD [3], both players have a certain chance to learn the current choice of the partner. Three cases are possible:

1. Player 1 sees the choice of Player 2 before making its own choice – with probability 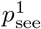.
2. Player 2 sees the choice of Player 1 before making its own choice – with probability 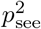.
3. Neither of players knows the choice of the partner, probability of this case is 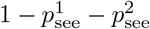.

Note that 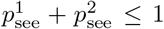 meaning that in each round only one of the players can see the partner’s choice. In the transparent iPD it is natural to assume that both players act on average at the same time [3]. Indeed, though both players are interested in the partner’s choice and would wait for the partner’s action, there is always a time constraint preventing players from waiting indefinitely. If this time constraint is explicit and known to both players, in most cases they are motivated to act just before the end of the time allowed for the action, thus nearly at the same time. Therefore for the rest of the paper we assume that 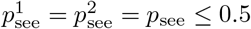 and call *p*_see_ the transparency level.

As it is usually done for the simultaneous iPD [5], we assume that players take into account outcomes of the previous game round, that is, use “memory-one strategies”. Then a strategy of a player in a transparent iPD is represented by a vector 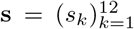, where *s_k_* are conditional probabilities to cooperate in 12 different situations. These depend on whether the player and the partner cooperated in the previous round, whether the player can see the current choice of the partner, and what the choice is if it is visible, see Table 1.

**Table 1.**
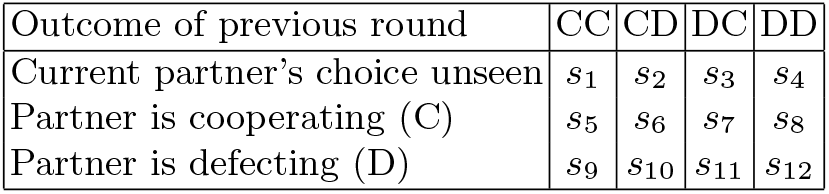
Representation of strategies in transparent games. Each strategy entry *s_i_* specifies probability of a player to cooperate depending on the outcome of previous round (the first action specifies the choice of the player, and the second the choice of the partner) and whether current partner’s choice is visible.

To find out what strategies are optimal depending on the transparency level p_see_, we use evolutionary simulations described in the following subsection.

### 2.2 Evolutionary dynamics of memory-one strategies in transparent games

To study evolutionary dynamics of the transparent iPD, we used the techniques developed for the simultaneous iPD in [4, 5] with minor adaptation to account for the different strategy representation in transparent games, see also [3].

Consider a population consisting of transparent-iPD players from n “species” defined by their strategies 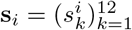 for *i* =1, 2,…, *n*. That is, any species i is a group of players sharing the same strategy s_*i*_, which they use when playing the iPD with a given transparency level *p*_see_ ∈ [0.0,0.5] against any partner. We assume that the population is infinitely large since a finite number of players results in complex stochastic effects [10], which we do not consider here. The population evolves in generations *t* = 1, 2,…. In each generation every player plays infinitely many rounds of the transparent iPD against a partner assigned randomly according to the current composition of the population. Species getting higher average payoff than others reproduce more effectively, and their relative frequency *x_i_*(*t*) in the population increases in the next generation. Note that 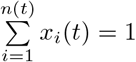 for any generation *t*.

Specifically, the evolutionary success of species *i* is encoded by its fitness *f_i_*(*t*), computed as the average payoff for a player from this species when playing against the current population:

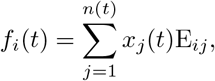

where E_*ij*_ is the expected payoff of species *i* playing against species *j*. If *f_i_*(*t*) is higher than the average fitness of the population 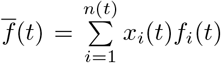, then *x_i_*(*t*) increases with time, otherwise *x_i_*(*t*) decreases and the species is dying out.

This evolutionary process is formalized by the replicator equation [5]:

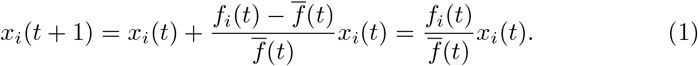

It remains to compute the expected payoff E_*ij*_ for a strategy 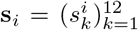 against 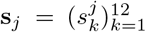. For this, consider Players 1 and 2 from species *i* and *j*, respectively. Since both players use memory-one strategies, their choices in every round of the game depend only on their mutual choices in the previous round. This allows to describe the game dynamics by a Markov chain with four states being the mutual choices of the two players (CC, CD, DC and DD), and transition matrix given by

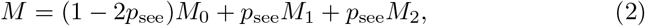

with matrices *M*_0_, *M*_1_ and *M*_2_ describing the cases when neither player sees the choice of the partner, Player 1 sees the choice of the partner before making own choice, and Player 2 sees the choice of the partner, respectively.

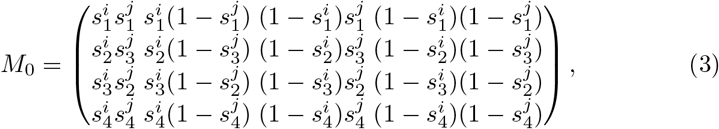

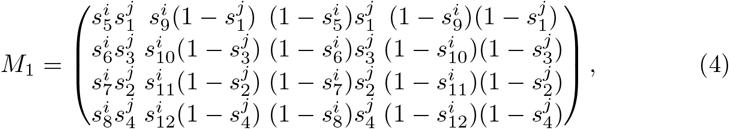

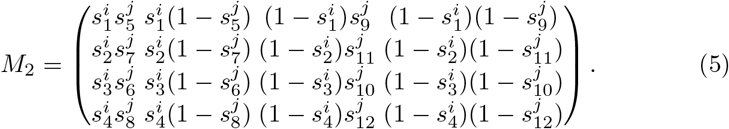

Then we can represent the expected payoff by the following formula

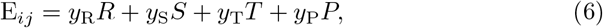

where *R, S, T, P* are the entries of the payoff matrix (*R* = 3, *S* = 0,*T* = 5, *P* =1 for the standard iPD, see Fig. 2), and *y*_R_,*y*_S_,*y*_T_,*y*_P_ represent the probabilities of getting to the states associated with the corresponding payoffs by playing s_*i*_ against S_*j*_. This vector is computed as a unique left-hand eigenvector of matrix *M* associated with eigenvalue one [5]:

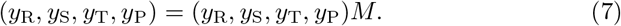

To guarantee the existence of the eigenvector (*y*_R_, *y*_s_, *y*_T_, *y*_P_), strategy entries for any species *i* should satisfy the following double inequality

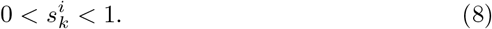

for all *k* =1, 2,…, 12. Therefore, it is common to introduce a minimal possible error ε in the strategies such that 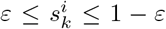. This also allows accounting for error-proneness of players (so-called “trembling hand” effect [4,11]).

Equation (6) allows to compute the expected payoff E_*ij*_ for all strategies *i,j* in the population. The dynamics of the population is entirely described by the matrix E. This dynamics is relatively simple for two species, when only four cases are possible [10]:

-*Dominance:* one species is unconditionally more fit than the other and replaces it in the population. Strategy *i* dominates *j* when E_*ii*_ > E_*ji*_ and E_*ij*_ > *E_jj_*.
-*Bistability:* either species can take over the whole population depending on the initial relative frequencies *x_i_*(1), *x_j_*(1) and the threshold *x*\*, given by the (non-stable) equilibrium frequency of species *i*

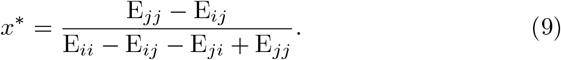
-If *x_i_*(1) > *x*\*, then species *i* takes over the population, otherwise species *j* wins. Bistability occurs when E_*ii*_ > E_*ji*_ and E_*ij*_ < E_*jj*_.
-*Coexistence:* the population is composed of two species with the asymptotic frequencies *x_i_* = *x*\*, *x_i_* = 1 – *x*\*, where *x*\* is a stable equilibrium given by (9). This case takes place when E_*ii*_ < E_*ji*_ and E_*ij*_ > E_*jj*_.
-*Finally, neutrality* takes place when E_*ii*_ = E_*ji*_ and E_*ij*_ = E_*jj*_, meaning that the frequencies of the species do not change.

We used the analysis of two-strategy dynamics for a pairwise comparison of strategies. However, already for a population consisting of *n* = 3 species, the dynamics can be rather complex, since there are 33 possible types of dynamics [12]. Analytic considerations for *n* > 3 are even more complicated [13] and are usually replaced by evolutionary simulations [5]. Details of our simulations are provided in Section 2.3.

### 2.3 Evolutionary simulations

For the evolutionary simulations we adopted methods suggested in [4,5]. We studied the populations of transparent iPD players for various transparency levels *p*_see_, and for each level we ran multiple evolutionary simulations as described below.

In each run, the population evolved as described in Section 2.2, thus the frequencies of species *x_i_*(*t*) changed according to (1). When *x_i_*(*t*) < χ, the species *i* was assumed to die out and was removed from the population; its fraction *x_i_*(*t*) was distributed among the remaining species proportional to their share in the population. We followed [4, 5] in taking χ = 0.001.

In each run a new species could enter the population with probability 0.01, thus new species emerged on average every 100 generations. Following [4], probabilities 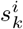 for the new species were randomly drawn from the distribution with U-shaped probability density function (10), favoring probability values around 0 and 1:

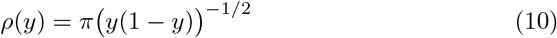

for *y* ∈ (0,1). We required 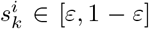 with *ε* = 0.001 as suggested in [4,5] to satisfy inequality (8). Initial frequencies of the introduced species were set to 1.1χ with χ = 0.001 [4].

The most important strategy entries, especially for low transparency, are s_1_, s_2_, s_3_, s_4_ (probability to cooperate when partner’s choice is unknown), since players use one of them with probability 1 – *p*_see_ > 0.5. Therefore these entries converge to optimal values quite fast and their values in evolutionary successful strategies are most precise, while convergence of other entries to optimal values may take longer. Since values of s_1_, s_2_, s_3_, s_4_ for evolutionary successful strategies were described in [3], here we limited their variability to allow faster convergence of the remaining entries s_5_,…, s_12_ to the optimal values. For this we rounded strategy entries s_1_, s_2_, s_3_, s_4_ so that they had the values from the set

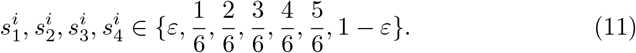

For the iPD with the payoff matrix shown in Figure 2 commonly discussed strategies are formed by the values from the set 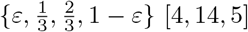, but in 11 we also consider intermediate values to achieve better discretization.

We carried out evolutionary simulations with random and with pre-defined initial compositions of the population. A random initial population consisted of *n*(1) = 5 species with equal frequencies *x*_1_(1) =…= *x*_5_(1) = 0.2 and random strategies. In this case we traced 10^9^ generations in each run. We also considered three pre-defined initial populations consisting of a single species with one of the following strategies:

-Win stay, lose shift (WSLS): s = (1,0,0,1; 1,0,0,1; 0, 0,0,0);
-Generous tit-for-tat (GTFT): 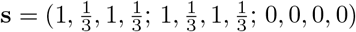;
-Leader-Follower (L-F):

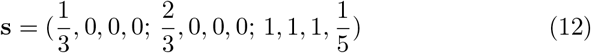

These strategies were selected as most evolutionary successful in simulations with random initial population for various transparency levels *p*_see_ (see Section 3 for details). For the pre-defined initial populations we traced 10^8^ generations in each run since it was not necessary to ensure stabilization of the population dynamics (which was the case for the random initial population). For each of the four initial compositions of the population described above, we performed 80 runs for each value of *p*_see_ = 0.0,0.1,…, 0.5. Additional 80 runs of simulations were performed for transparency levels *p*_see_ = 0.26,0.28,…, 0.50, for the initial WSLS and L-F populations.

### 2.4 Measuring the strategies’ evolutionary success

To quantify the evolutionary success of a strategy we computed the average of its relative frequency *x_i_*(*t*) across all generations t and across all runs. Since the strategies in the evolutionary simulations were generated randomly, some of the observed strategies might not entirely converge to the theoretical optimum. Therefore, we used a coarse-grained description of strategies as suggested in [3] with the following notation: symbol 0 for 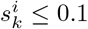, symbol 1 for 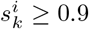, symbol * is used as a wildcard character to denote an arbitrary probability. We characterized as Always Defect (AllD) all strategies encoded by (0000;**00;**00), meaning that the probability to cooperate when not seeing partner’s choice or after defecting is below 0.1, and other behavior is not specified. Similarly, the generalized representations of other strategies were as follows

-Win stay, lose shift (WSLS): (100*c*;1***;****) with *c* ≥ 2/3;
-Tit-for-tat (TFT): (1010;1***;****);
-Generous tit-for-tat (GTFT): (1*a*1*c*;1***;****), where 0.1 < *a, c* < 0.9;
-Generous WSLS (GWSLS): (1*abc*;1***;****), where *c* ≥ 2/3, *a,b* < 2/3 and either *a* ≥ 0.1 or *c* > 0.1;
-Firm-but-fair (FbF): (101*c*;1***;****), where 0.1 < *c* < 0.9;
-Leader-Follower (L-F): (*00*c*;****;*11*d*), where *c* < 1/3 and *d* < 2/3.

## 3 Results

Frequencies of strategies for various initial compositions of the population are presented in Fig. 3. In all cases, the fraction of the described strategies drops down for *p*_see_ > 0.3. This happens due to the fact that for higher transparencies population dynamics often enters a “chaotic” mode, where many transient strategies replace each other in a rapid succession [3]. Each of these transient strategies has a low relative frequency, while taken together they constitute a considerable fraction of the population, especially for high *p*_see_.

**Fig. 3.**
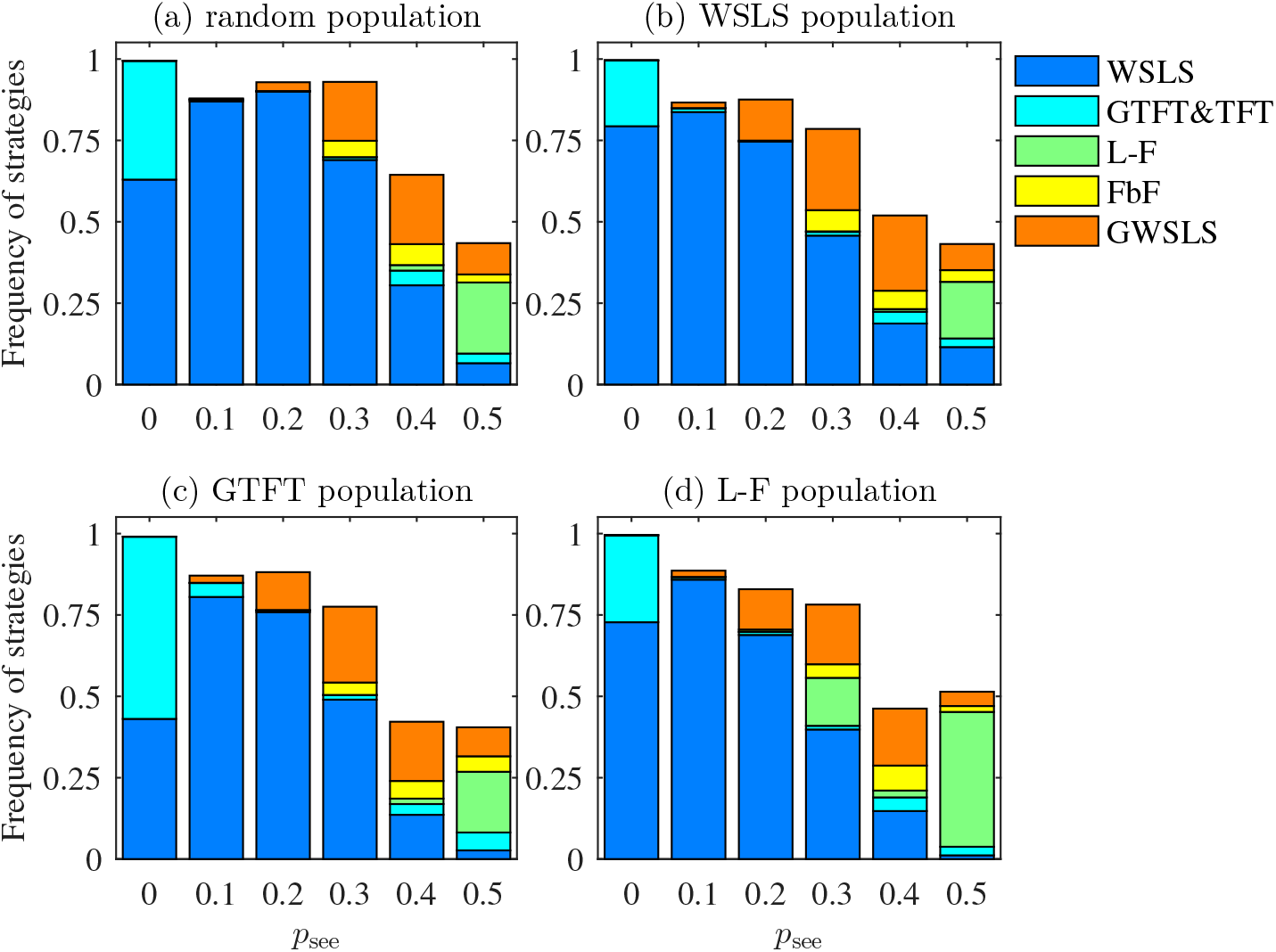
Average relative frequencies of strategies occurred in the population for its various initial compositions: (a) random; constituted by (b) WSLS, (c) GTFT and (d) L-F strategies. Other strategies are transient (i.e. persist in the population only for a relatively short time). Results for all initial populations are quite similar, yet a few differences are noteworthy. First, the frequencies of WSLS and GTFT depend on the initial composition of the population for *p*_see_ = 0, but not for higher transparency. When one of these strategies was the initial strategy of the population, it is much more abundant than for the random initial composition. Second, when L-F is the initial strategy of the population, it is has considerably higher frequency for *p*_see_ = 0.3 and *p*_see_ = 0.5, but not for *p*_see_ = 0.4. Since for *p*_see_ = 0.3 L-F has a noticeable frequency only when forming the initial population, for this transparency L-F can persist but cannot take over the population when starting with a low frequency (see Fig. 4b). Note that such non-monotonicity in strategy dynamics is not altogether unexpected, since the dynamics depends non-linearly on the transparency level [3].

Regardless of the initial population composition, “Win-stay, lose-shift” (WSLS) strategy is a clear winner for 0 < *p*_see_ < 0.4. The theoretically optimal form of this strategy is (1001;1001;0000), however in simulations it may appear as (1001;1**1;000*), where the entries marked by the wildcard character * can take a value from 0 to 1 (see Fig. 5).

**Fig. 4.**
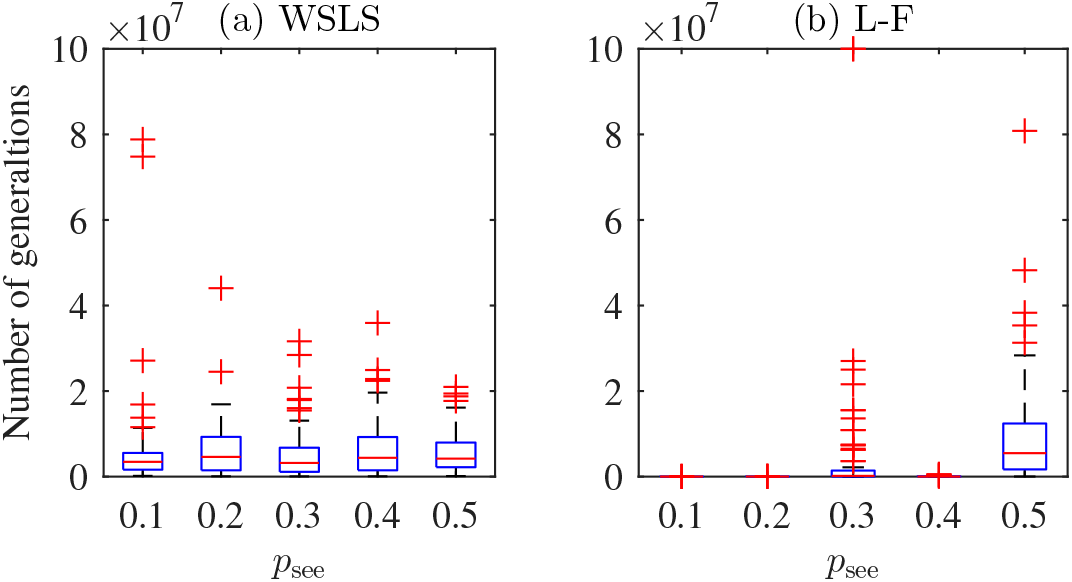
Number of generations for that initial strategy of the population remains the most frequent for the initial population constituted by (a) WSLS and (b) L-F players. The central mark indicates the median, and the bottom and top edges of the box indicate the 25th and 75th percentiles, respectively. The whiskers extend to the most extreme data points not considered outliers, and the outliers are plotted individually using the + symbol. The higher the number of generations, the longer the initial strategy persists in the population. While persistence of WSLS is relatively stable, L-F can persist only for *p_see_* = 0.3 and *p_see_* = 0.5.

**Fig. 5.**
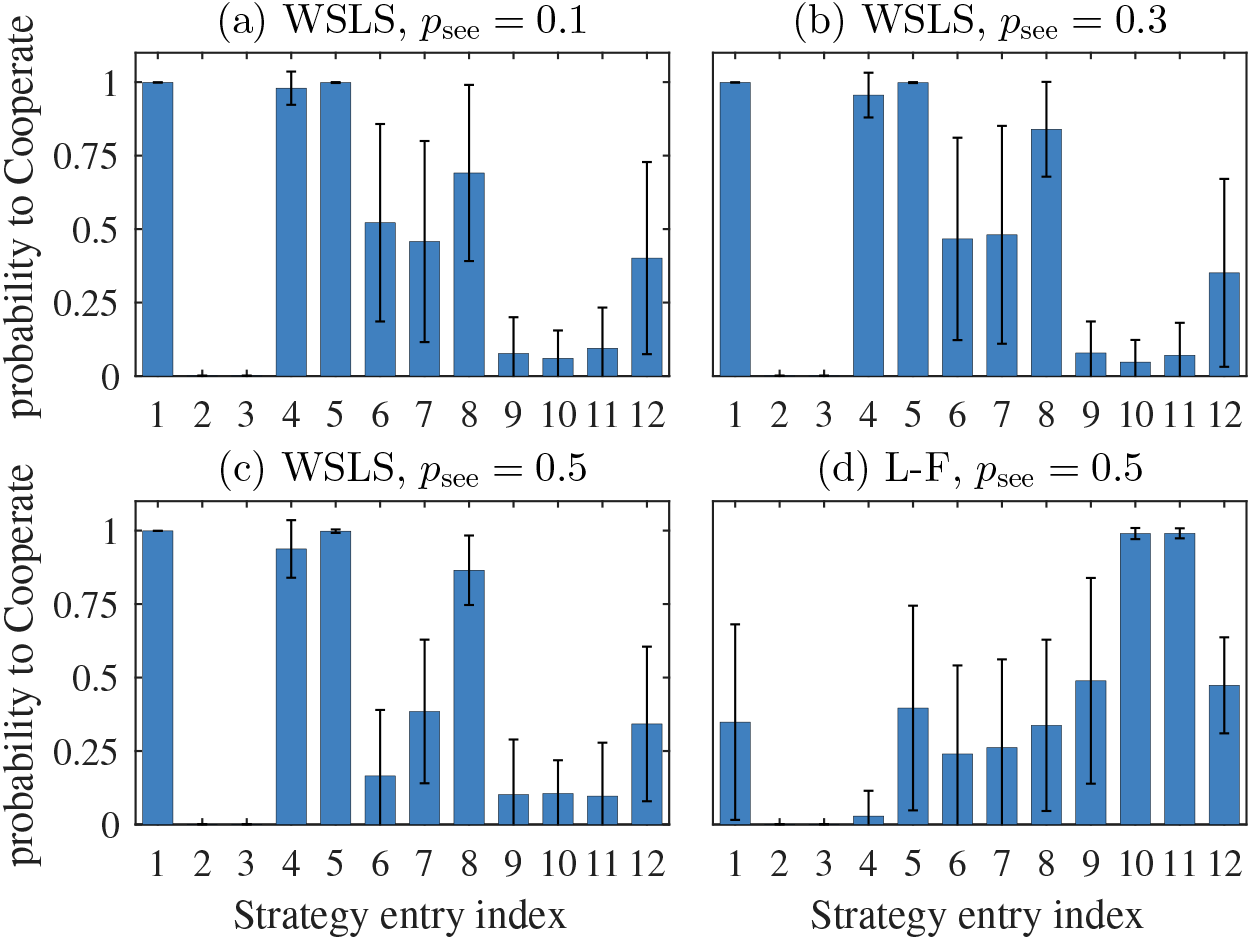
Average strategy profiles with standard deviations for the winning strategies in a transparent iPD: (a) for WSLS at *p*_see_ = 0.1, (b) *p*_see_ = 0.3, (c) *p*_see_ = 0.5 and for L-F at *p*_see_ = 0.5 (d). Each strategy profile is represented by 12 entries characterizing probability of a player to cooperate. Namely, *s*_1_,…,*s*_4_ are probabilities to cooperate when current partner’s choice is unknown and the outcome of the previous round was “both cooperated”, “self cooperated, partner defected”, “self defected, partner cooperated” and “both defected”, respectively. Similarly, *s*_5_,…, *s*_8_ and *s*_9_,…, *s*_12_ are probabilities of a player to cooperate seeing that current partner’s choice is to cooperate and to defect, respectively. Standard deviations show the difference in variability for various entries. For instance in (a) some strategy entries are constant *s*_1_, *s*_2_, *s*_3_, *s*_5_, some vary only slightly *s*_4_, *s*_9_, *s*_1_o, *s*_11_, and other entries are almost random *s*_6_, *s*_7_, *s*_8_, *s*_12_.

For *p*_see_ ≥ 0.1 WSLS can be only replaced in the population by a strategy (1001;*d*\*\*\*;*00*) with 0 < d < 1 (Fig. 6a-c). In [3] this strategy was called treacherous WSLS since it behaves like WSLS when not seeing the choice of the partner, and defects when seeing that partner cooperates. Treacherous WSLS has low payoff when playing with itself and is easily replaced by other strategies, but it dominates WSLS for any *p*_see_ > 0.

**Fig. 6.**
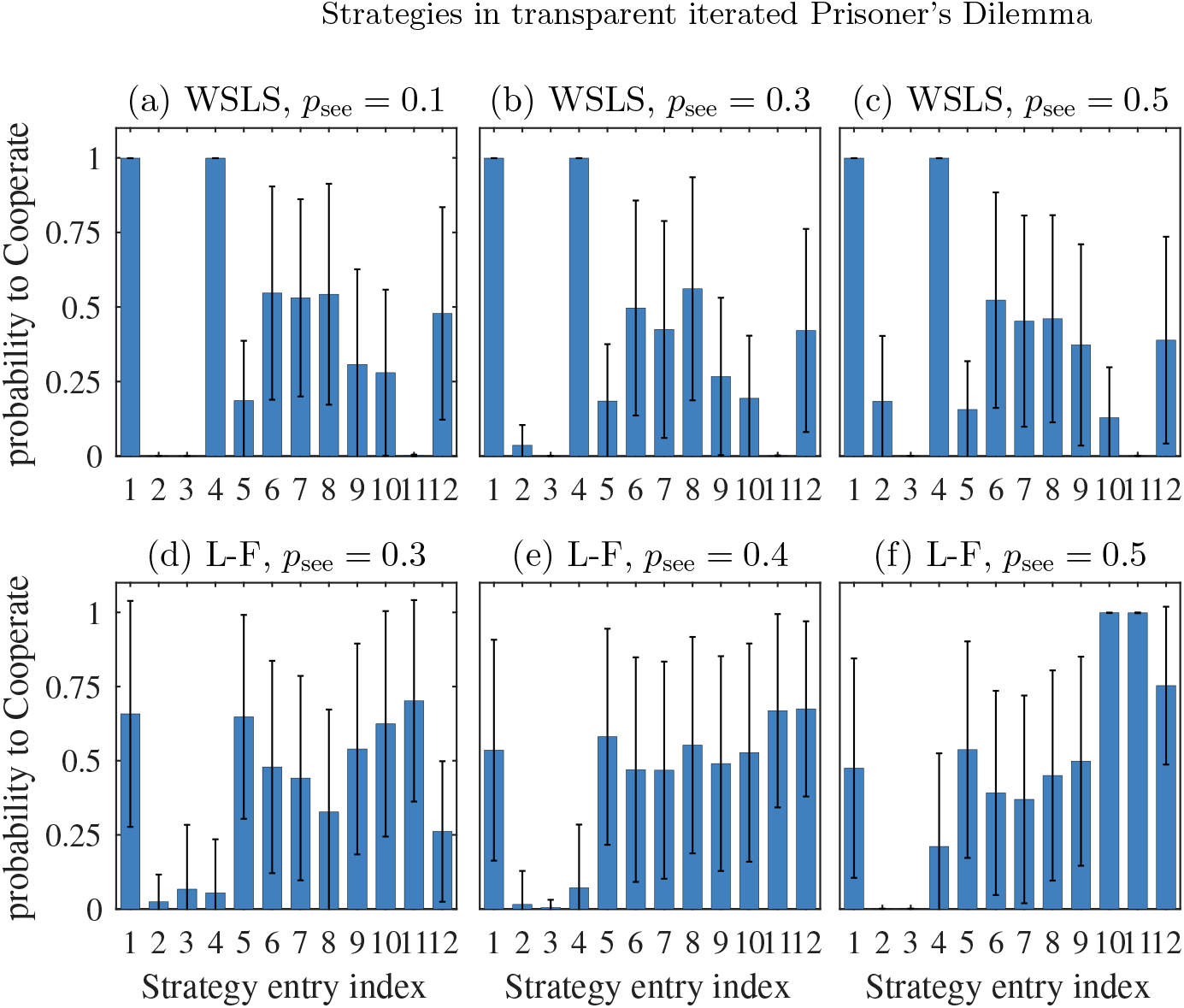
Average strategy profiles with standard deviations for the strategies replacing initial strategies in simulations with pre-defined initial population: (a)-(c) strategies replacing WSLS for *p*_see_ = 0.1, *p_see_* = 0.3 and *p*_see_ = 0.5, respectively; (d)-(f) strategies replacing L-F for *p*_see_ = 0.3, *p*_see_ = 0.4 and *p*_see_ = 0.5, respectively.

The predominance of WSLS as the single frequent non-transient strategy can be challenged only for *p*_see_ ≈ 0 (minimal transparency, nearly-simultaneous iPD) by GTFT and for *p*_see_ > 0.4 (maximal transparency, nearly-sequential iPD) by L-F. Below we discuss both these cases in detail.

For minimal transparency the WSLS frequency slightly drops (Fig. 3a) since in this case population of WSLS players can be invaded by AllD. Let s_1_ = (1 – *ε, ε, ε*, 1 – *ε*; 1 – *ε, ε, ε*, 1 – *ε; ε, ε, ε, ε*) (WSLS), s_2_ = (*ε, ε, ε, ε; ε, ε, ε, ε; ε, ε, ε, ε*) (AllD). Estimating matrix of expected payoff by (6), we see that 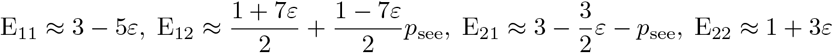.

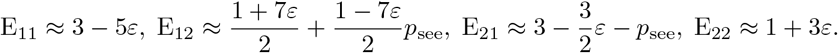

Thus for *p*_see_ ≤ 3.5ε WSLS is dominated by AllD and for *p*_see_ > 3.5ε the two strategies are bistable. AllD invades WSLS in the former case unconditionally and in the latter case if its equilibrium frequency given by (9) is sufficiently low. Namely, if AllD with initial fraction 1.1χ is introduced to the population playing WSLS, for 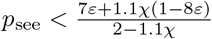 AllD invades the population. In this case, unconditional defectors can take over the population for a considerable number of generations, and strategies different from WSLS (TFT and GTFT) should be introduced to the population to restore cooperation. These strategies can resist WSLS invasion, therefore once WSLS dies out it may take many generations to re-establish itself.

In our simulations, *ε* = 0.001, *χ* = 0.001, so WSLS can resist AllD-invasion for *p*_see_ > 0.0041. In this case WSLS is dominated only by treacherous WSLS. This weak transient strategy is easily replaced by others, which allows WSLS to reappear in the population relatively quickly.

The decrease of WSLS frequency for high transparency (Fig. 3, 7) is caused by two factors. First, as one can see from Fig. 5, the higher *p*_see_ the more precisely WSLS should correspond to its theoretically optimal profile. Indeed, for *p*_see_ = 0.1 most entries of WSLS have relatively high standard deviations, meaning that in this case a successful strategy can follow the WSLS principle in a rather general fashion. Meanwhile, for *p*_see_ = 0.5 most entries have low variability, meaning that successful WSLS variants can have only slight deviations from the optimal profile. The closer a strategy must be to the theoretical WSLS profile in order to be successful, the lower the probability that such a strategy is by chance introduced in the population, which results in lower frequency of WSLS for higher *p*_see_. Note that the frequency of WSLS would decrease even if we understand WSLS in the most general sense and consider GWSLS as a part of it (Fig. 8). Second, the higher *p*_see_ is, the faster and easier the treacherous WSLS takes over the population, since it has higher chances to defect WSLS. This also means that a strategy that only roughly resembles the treacherous WSLS profile can be successful against WSLS for high *p*_see_ (see Fig. 6a-c).

**Fig. 7.**
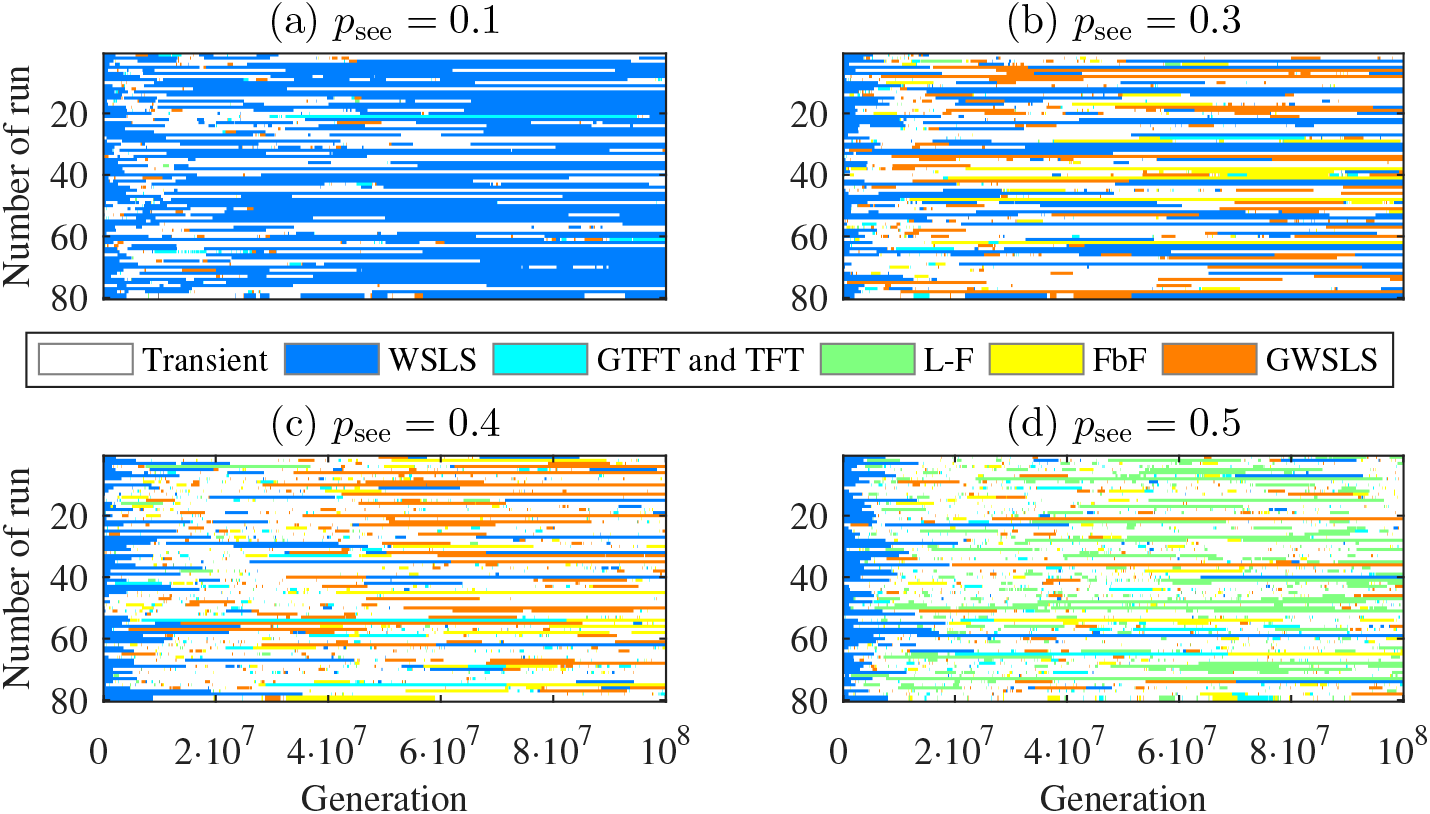
Dynamics of the most frequent strategy of the population in various runs for initial WSLS-population. (a) For *p*_see_ = 0.1 WSLS is predominant in most runs. (b) For *p*_see_ = 0.3 WSLS frequency wanes, while other TFT, L-F, FbF and GWSLS become more abundant. Fraction of various transient strategies also increases and (c) for *p*_see_ = 0. 4 they become most frequent. (d) For *p*_see_ = 0.5 WSLS rarely reestablishes itself and the only frequent strategy is L-F. Population mostly is in a chaotic mode with transient strategies replacing each other.

**Fig. 8.**
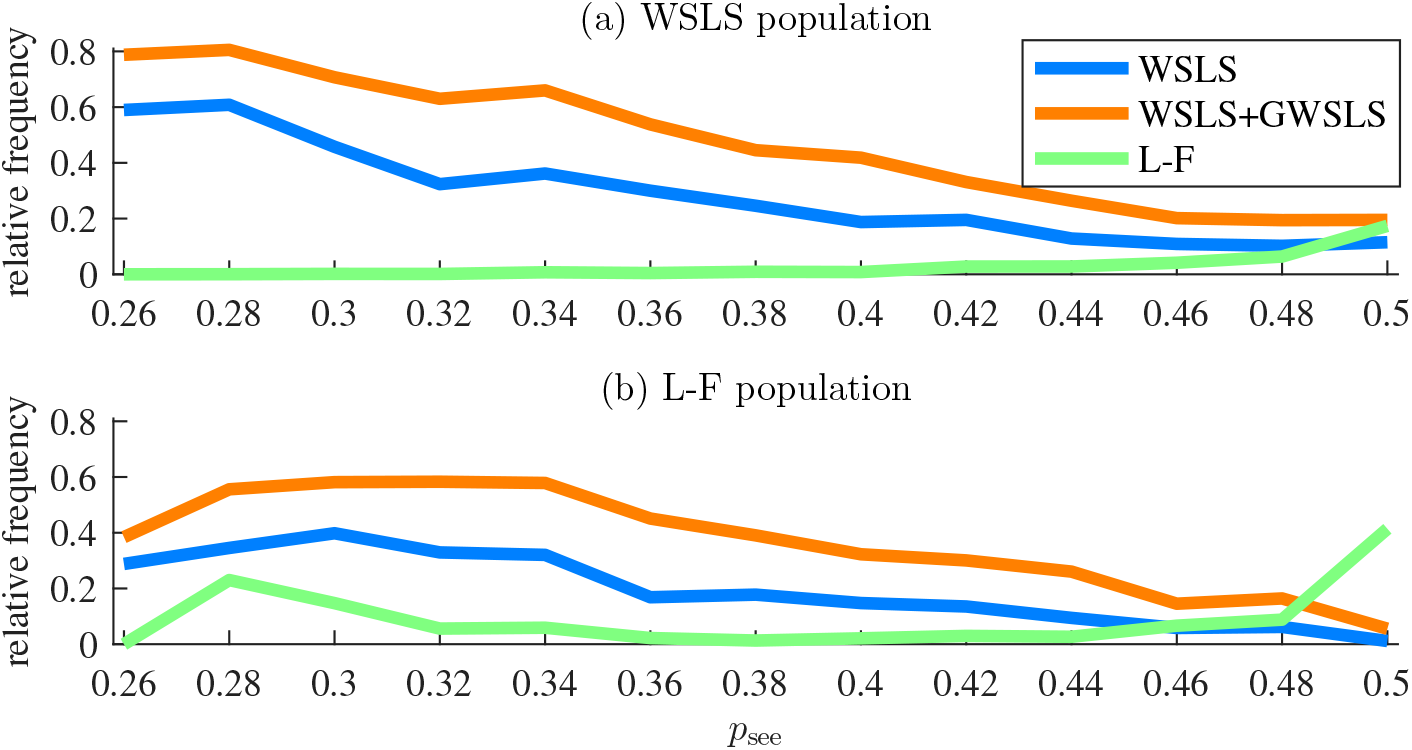
Average relative frequencies of WSLS, WSLS with GWSLS, and L-F strategies for the initial populations constituted by (a) WSLS player and (b) L-F players. Note that in both cases WSLS frequency decreases considerably for *p*_see_ > 0.35.

It remains to explain the success of L-F for high transparency. Consider the effect of *p*_see_ on the frequencies of WSLS and L-F in more detail. Fig. 8 shows how these frequencies vary for *p*_see_ ∈ [0.26,0.50] taken with a step of 0.02. for the initial population consisting either of WSLS or L-F players. Note that L-F is only successful for maximal transparency *p*_see_ = 0.5 (Fig. 8a), although for some transparency levels *p*_see_ < 0.5, L-F can remain in the population once introduced (Fig. 8b).

The success of L-F for maximal transparency can be partially explained by the fact that a variant of this strategy given by equation (12) can be replaced in the population only by other variants of the L-F strategy (Fig. 6d). Although these variants are not evolutionary stable and are eventually replaced by other strategies (Fig. 3d), this indicates a good evolutionary potential of L-F for *p*_see_ = 0.5. One particularly interesting modification of L-F is a strategy with a profile (1001;1001;1111). As it combines the features of L-F and WSLS we term it “WSLS-like L-F”. This strategy dominates normal L-F and easily replaces it in the population. But contrary to the normal L-F, WSLS-like L-F is highly unstable since it is dominated by many strategies including WSLS, treacherous WSLS and AllD. Therefore, WSLS-like L-F stays in population just for a few generations and then is replaced by other strategies (similar to the treacherous WSLS).

## 4 Conclusion

Here we aimed to explain the success of various strategies in the transparent iterated Prisoner’s dilemma [3]. Our main findings are:

-Win stay, lose shift (WSLS) and Generous tit-for-tat (GTFT) are two predominant strategies for *p*_see_ ≈ 0. This case of low transparency is very close to the classical simultaneous Prisoner’s dilemma in which agents make choices without knowing the respective choice of the other agent.
-For most non-zero transparency levels (0 < *p*_see_ < 0.5), WSLS is the only effective memory-one strategy. However, WSLS is not evolutionary stable, since it can be counteracted by its modified version, termed “treacherous WSLS”, which defects when seeing the partner’s choice.
-For *p*_see_ ≈ 0.5 (maximal transparency) a second strategy, Leader-Follower (L-F), becomes predominant. When two players use this strategy, one of them defects and the other cooperates, leading to a kind of “turn-taking” behavior. However, L-F is also not evolutionary stable since it evolves towards highly unstable WSLS-like variants.

Fig. 9 summarizes our results schematically representing the strategy dynamics for various transparency levels *p*_see_.

**Fig. 9.**
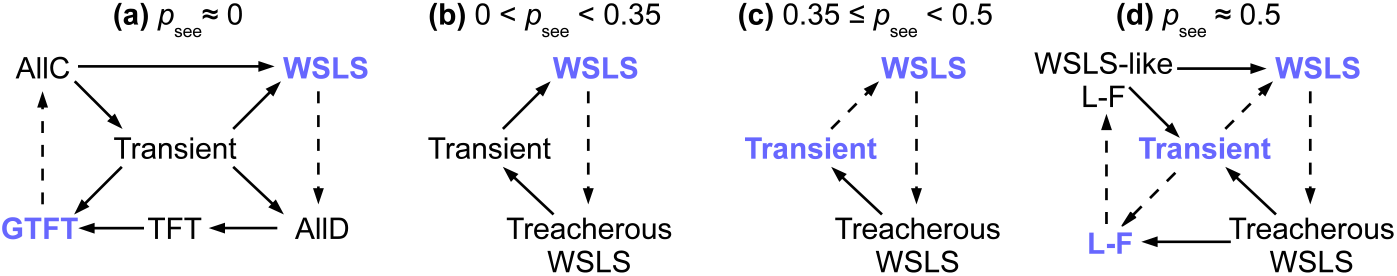
Strategy dynamics for various values of *p*_see_. Solid and dashed arrows indicate likely and less likely directions of dynamics, respectively. Strategies that can persist in the population for high number of generations (compared to other strategies) are shown in bold blue font. We group all strategies that cannot persist in the population and do not have special importance for the dynamics as “transient”. Four cases of dynamics can be distinguished. (a) For *p*_see_ ≈ 0 (for *p*_see_ < 0.004 in our simulations) the game resembles the classic simultaneous iPD. In this case strategy dynamics is relatively complex [5]. The population oscillates between two predominant strategies, WSLS and GTFT, and the transient states are relatively short. (b) As we increase *p*_see_, two important changes take place. First, GTFT becomes ineffective against other strategies, including WSLS, L-F and AllC. Second, for *p*_see_ > 0.004 AllD cannot invade a population of WSLS players, and the only strategy that can “dethrone” WSLS is treacherous WSLS. These changes result in greatly simplified dynamics with WSLS being the only non-transient strategy. It is occasionally invaded by treacherous WSLS, which controls the population only for a short time and is then replaced by other strategies. (c) For high transparency (0.35 < *p*_see_ < 0.5, with threshold value 0.35 selected based on dynamics of strategy frequencies shown in Figure 8), WSLS is still predominant, but it becomes difficult for this strategy to take over the population. Therefore, the population spends most of the time in a “chaotic” state with various transient strategies quickly replacing each other. (d) Finally, for maximal transparency (*p*_see_ ≈ 0.5), L-F becomes the second predominant strategy. Yet, L-F does not have stable control of the population since some modifications of L-F can be invaded by non-L-F strategies. As a result, transient strategies still control the population for most of the time.

Overall our results provide an important extension to previous studies that have used evolutionary simulations to suggest and test strategies in classic simultaneous and sequential games. By concentrating on transparent games frequently encountered in real life but rarely investigated, we provide a partial explanation for the fact that the strategies developed for the simultaneous iPD are rarely observed in nature [6, 7], where high levels of transparency are common [1].

## Acknowledgments

We acknowledge funding from the Ministry for Science and Education of Lower Saxony and the Volkswagen Foundation through the program “Niederschsisches Vorab”. Additional support was provided by the Leibniz Association through funding for the Leibniz ScienceCampus Primate Cognition and the Max Planck Society.

